# A review of online learning strategies for biological NMR techniques a survey of educators

**DOI:** 10.1101/2023.01.13.523927

**Authors:** Marie M Phelan

## Abstract

Disruption to teaching over the pandemic has led to a range of online and digital solutions. Given the utility of software and remote platforms for NMR teaching a UK-based survey for NMR educators in undergraduate and postgraduate teaching was rolled out Spring 2022 in order to establish what online tools and techniques are employed to teach NMR to biological sciences and appraise their relative efficacy. Based on the survey outcomes an overview of the breadth of methods employed as well as their perceived effectiveness is presented with a list of recommendations for common and successful strategies for online (and hybrid) teaching of NMR to the life sciences.

## 1 Introduction

### 1.1 Aim & Objectives

The range of approaches to teach NMR at Degree and post-graduate level varies widely between higher education and discipline. Remote teaching pre-pandemic included virtual reality and online case studies with majority of learning resources focussed on chemistry applications.

In 2020 and 2021 need for remote educational tools became essential for all higher education institutes and this necessity lead to a renewed interest in online and computer based resources for teaching and learning NMR. Due to a variety of applications available and the sparsity of knowledge on effective strategies for remote teaching a survey of biological NMR educators was deployed. The aim of this project is to establish what online tools and techniques are employed to teach NMR to biological sciences and appraise their relative efficacy.

Objective 1: survey the UK NMR community to establish what methods were employed (and their perceived effectiveness relative to pre-pandemic teaching)

Objective 2: Analyse the results of the survey (with follow-up interviews where invited by the educator - optional question in survey) to determine what approaches were most suitable (and what resources were most heavily relied upon).

Objective 3: Based on objective 2 prepare a list of recommendations for common and successful strategies for online teaching of NMR to life sciences. (include limitations to each and mitigations where appropriate).

### 1.2 Background

A review of education literature for NMR heavily features articles from two education-focussed journals which featured NMR-teaching articles; “Journal of Chemical Education” (established 1924 ISSN:1938-1328) and “Biochemistry and Molecular Biology Education” (established 1972 ISSN:1539-3429). Other research tools employed for teaching are present throughout the literature were reviewed (such as in Protein Science special issue). Articles widely varied in aims (foundation-level problem solving and debate [Stowe & Cooper 2019] through to ways to appropriately describe spin evolution and pulse sequences [Vega-Hernandez & Antuch 2014, Williamson 2019]) and scope. Both journals scope includes innovations, pedagogic approaches, skill-building lab-experiments, software for educational use, and review of multimedia material.

The education literature includes practical examples and data to work through as group problems [Graham *et al* 2016] along with lab and group activities [Rocks & Stockland 2020]. Innovations such as virtual reality software for molecular modelling [Gandhi *et al* 2021], 3D printing of 2D spectra [Jones *et al* 2021] even iPad interactive software [Li *et al* 2014]. Artificial intelligence simulations [Rzepa *et al* 2021] spin-simulation [Boldt 2011, Sengupta 2021] and real-time monitoring [Zientek *et al* 2014] are all tools proposed by the chemical NMR community to leverage online capabilities to more descriptive and illustrative teaching resources.

In 2020 and 2021 need for remote educational tools became essential for all higher education institutes and this necessity led to a renewed interest in online and computer-based resources for teaching and learning NMR. An emergency shift to online teaching does not represent best practice and may be limited in resources available [Abou-Khalil *et al* 2021]. However, reflections from approaches success and limitations of teaching in the period may identify tools and approaches that would benefit teaching and learning to biological NMR. Problems with passive learning or lecture based teaching in sciences, technology, engineering and mathematics (STEM) are well recognised [Cooper *et al* 2015]. STEM disciplines in higher education favour a major component of inquiry based teaching, in essence aligning with a constructivist teaching philosophy through action, reflection and construction [Piaget 1964, Lakatos 1970]. On the other hand for teaching fundamental concepts within science a cognitive approach of comprehension, memory and application is also well aligned [Piaget 1964]. Therefore, this work focusses on remote strategies to facilitate ‘active learning’, ‘construction’ and ‘application’ in biological NMR education.

### 2.1 Methodology

Remote evidence based teaching of NMR requires resources specific to NMR-topic, resources specific to hosting online classrooms (tele-conferencing software such as teams, zoom, skype) and interactive tools that facilitate or attempt to emulate approaches taken for granted from an in person teaching and learning experience [Pilkington & Hanif 2021, Lee & Yeong 2020]. As the recent years have somewhat required all teaching to be conducted at least partially online are view of ubiquitous teleconferencing software is not the focus of this survey (this will be no doubt be appraised extensively by pedagogic and andragogic practitioners in the coming years).

Instead, this survey is focussed on NMR-based education and as such aims to explore effective approaches to teaching and learning specific to biological NMR. However, while there is a focus on NMR and NMR-adjacent applications, the impact of group work and student lead discussions to effective teaching should not be overlooked and a range of online teaching and learning approaches were suggested to facilitate this aspect of the NMR-learning experience.

Where possible it was also important to challenge the preconception that pandemic teaching was by necessity less organised and less valuable to learning outcomes than pre-pandemic. Therefore, an attempt to compare outcomes was also included. Post-graduate and PhD teaching is not assessed granularly and as such may lead to limitations on how the change in teaching were valued. Therefore, follow-up discussions were employed (and supplemented with case studies).

### 2.2 Survey Design

#### Scope

The survey was designed in four sections – the first to establish the demographics of the participants (their location, teaching requirements, learners level and topics covered within biological NMR). Secondly the use of online and computational based software employed during remote teaching. The third section concerned measures of effectiveness or assessing success (requesting participants compare completion-rates, pass-rates and grade distribution with respect to in-person learning). Finally the participants were required to self-reflect on any benefits and limits of the remote environment before being invited to (optionally) participate in follow-up discussions

#### Ethics involved

The responses to the survey were completely anonymous unless the participant volunteered for follow-up discussion on aspects of their teaching practice. All disclosures in follow-up discussion have been anonymised. All data was held securely and anonymously, and only accessible to the author at the University of Liverpool. The survey and follow-up discussions were covered by the University of Liverpool ethics application 5402.

#### Rollout

The survey was deployed in spring 2022 and open until 30^th^ July 2022. The original period for interaction was extended due to time pressures both within the community and the author.

#### Analysis

Closed questions were ranked according to number of participants selecting them with VENN diagrams capped at the four most common responses. Open questions in the survey were categorised through content analysis into categories of learning; facilitating; sharing/collaborating; practicing; assessing.

## 3 Results

### 3.1 Survey statistics

The results of the survey explore the tools employed in the remote classroom and the reflections from educators as to what remote approaches brought and subtracted from the learning experience.

The responses were spilt 8:13 between UK and the rest of world with educators teaching mainly at masters and PhD level the majority but not all taught biochemistry (with remaining responders teaching chemistry), there were educators teaching across both disciplines as well as some also teaching in biology. The range of disciplines taught within the cohort was mainly NMR theory (16 at basic level including 3 at advanced level) and/or protein structure (Figure 1).

**Figure1.**
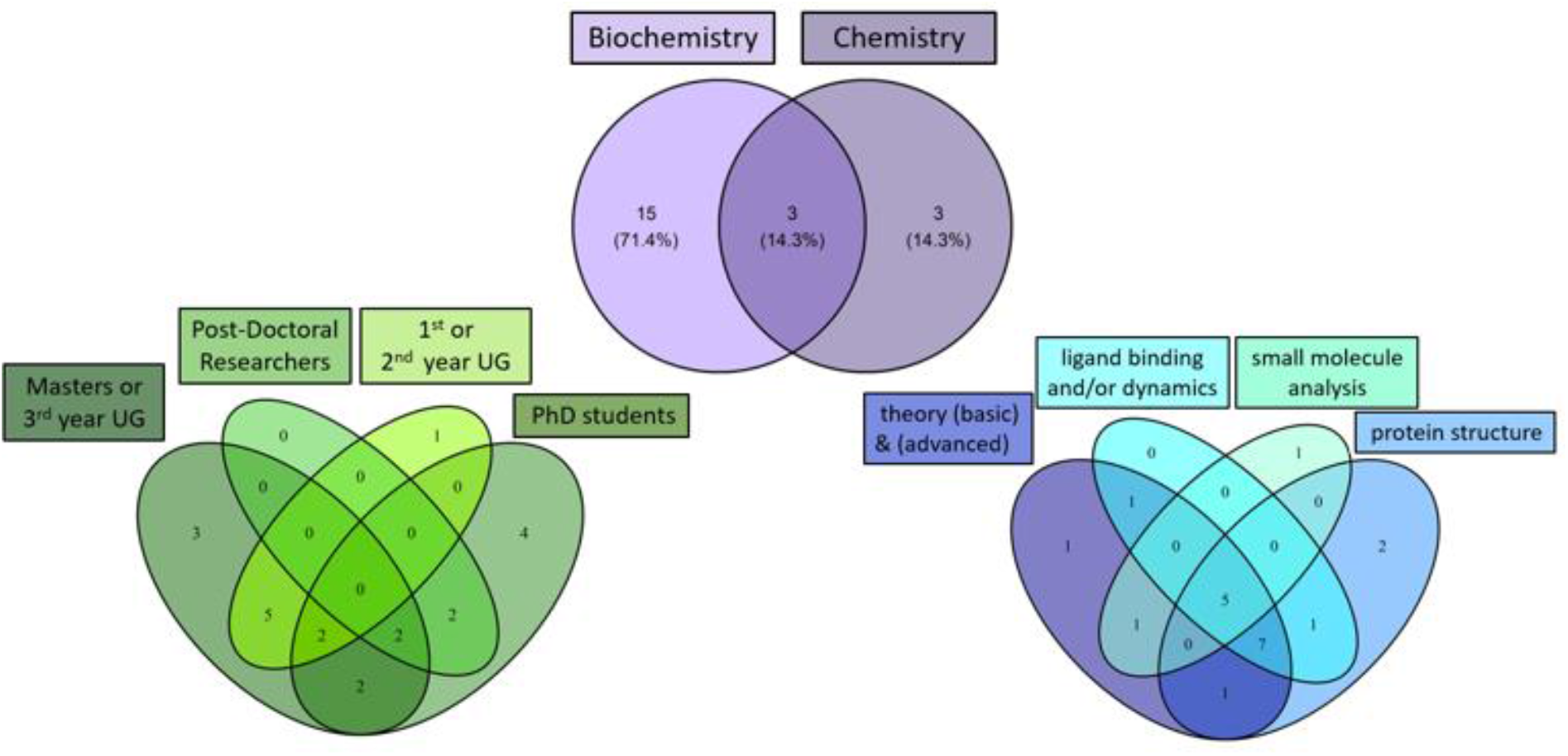
Demographic of Educators completing the Survey: top: disciplines taught, left: year of study taught, right: aspect of NMR taught.

A range of methods for delivery of remote teaching was employed with most favouring synchronous lectures (>90%) and at least half delivering problem solving workshops for spectral assignment (Table 1). employed by the majority (>76%, i.e. every respondent teaching protein structure). Over half of respondents (>61%) did not use online databases or prediction software for teaching and learning (Table 3).

**Table 1:** Methods for delivery of remote teaching.

| Method for Delivery | n | % |
| --- | --- | --- |
| synchronous lectures (slides and voice) | 19 | 90.4 |
| problem solving workshops - spectra assignment | 11 | 52.3 |
| asynchronous lectures (slides and voice) | 6 | 28.5 |
| small group tasks | 6 | 28.5 |
| polls, multiple choice quizzes etc. | 6 | 28.5 |
| video demonstrations | 3 | 14.2 |
| live practical demonstrations | 1 | 4.8 |
| 1-2-1 teaching | 1 | 4.8 |

While remote teaching of NMR was provided by every survey responder almost half reported no use of processing or assignment software (Table 2). Conversely Pymol atomic visualisation/modelling software was

**Table 2:**
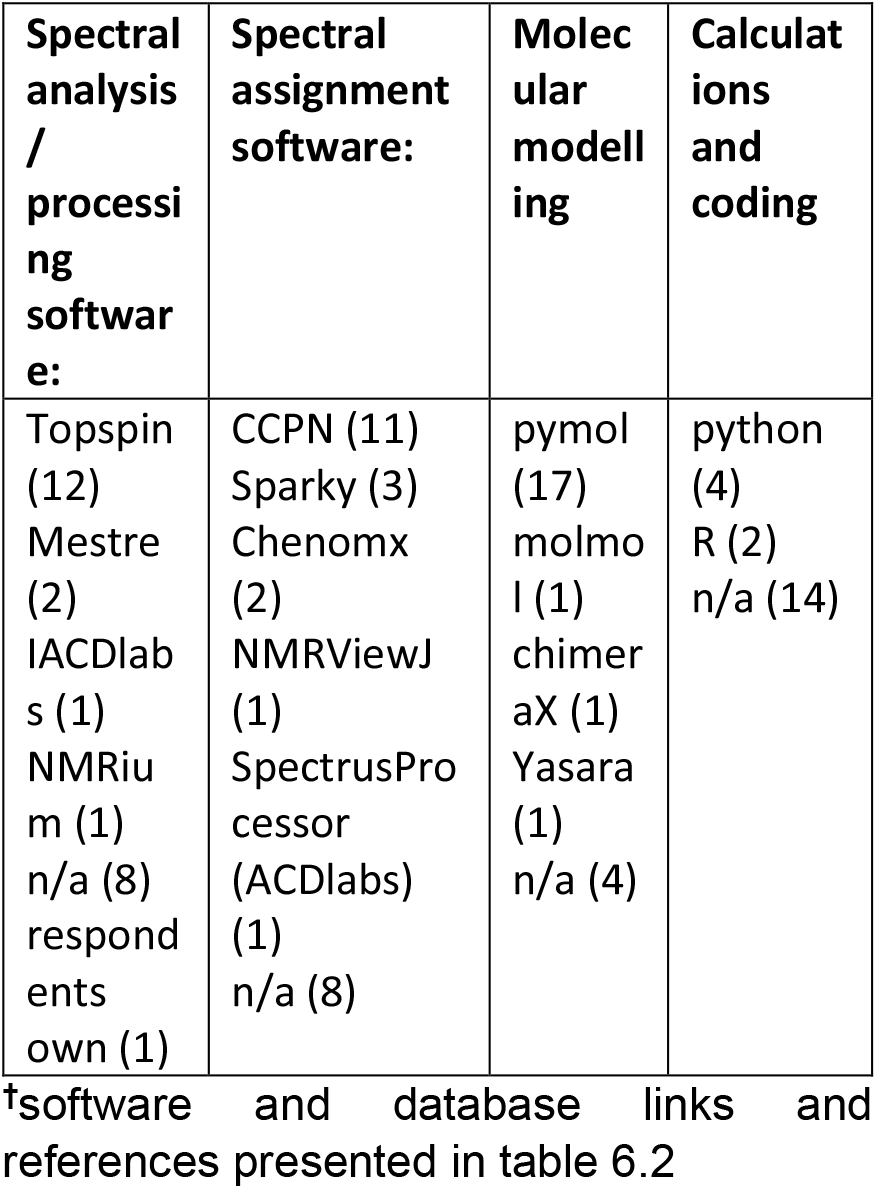
Software employed in the online learning environment for specific NMR aides^†^.

**Table 3:** Databases and Software employed in the online learning environment^†^.

| Databases | Prediction | Other |
| --- | --- | --- |
| BMRB (4) | NMRDB (2) | quizacademy (2) |
| SDBS (2) | ChemDraw (1) | moodle (H5P plugin) |
| EBI (2) | ACDlabs (1) | NMRbox |
| n/a (13) | n/a (13) | Nmrprocflow |
|  |  | PDB |
|  |  | respondents own |

<sup>†</sup>software and database links and references presented in table 6.2

### 3.2 Survey Educators measures for success

In order to establish whether educators experiences lead to effective teaching and learning strategies. As NMR in the biological sciences is often taught post-graduate there is a distinct split between formally graded outcomes [QAA] and developing professional practice [UKPSF; V4]. Furthermore, due to the range of assessment strategies within the survey cohort care is required not to over-interpret (Figure 2).

**Figure 2:**
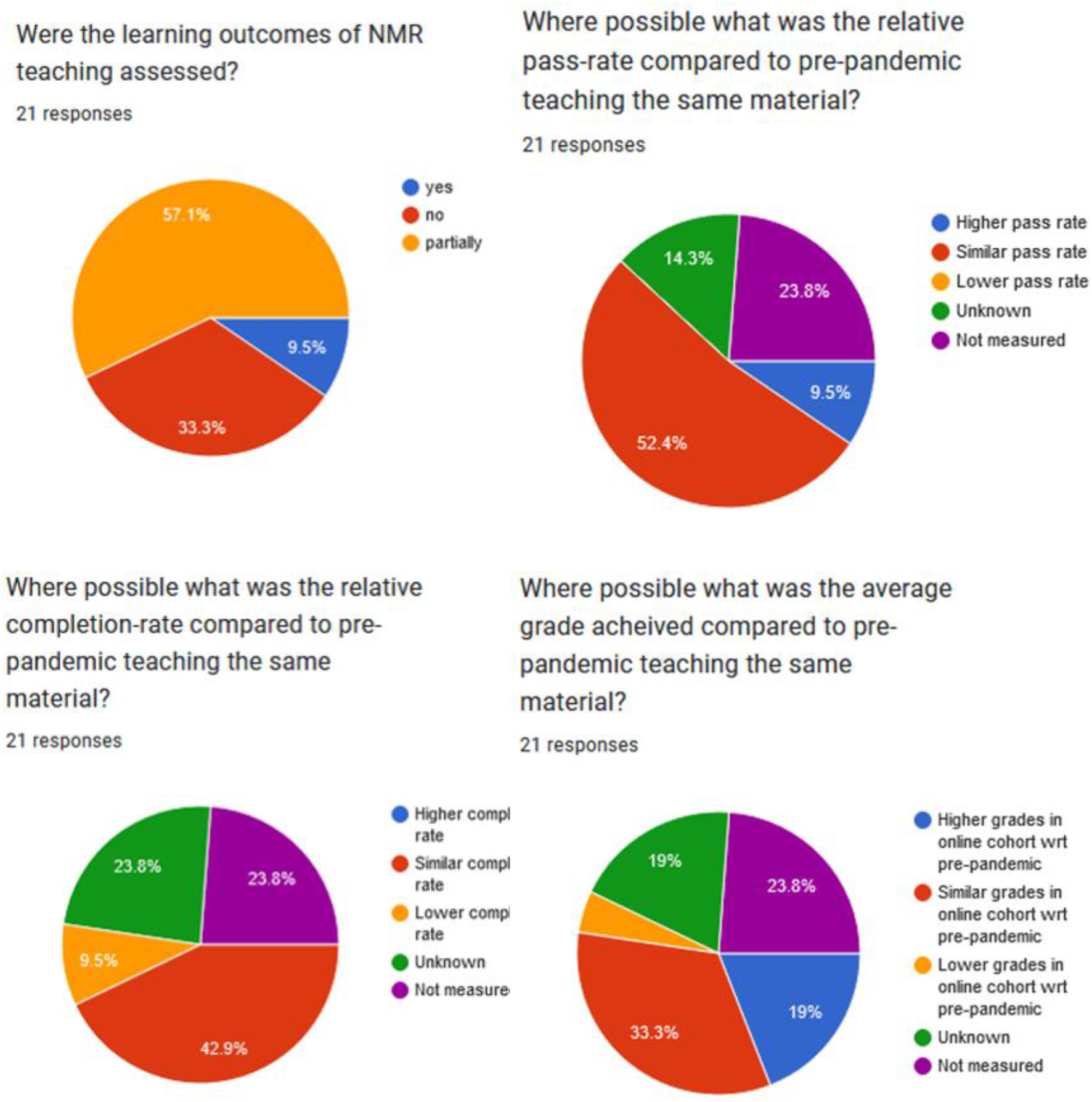
Measures of success. Participants were asked to compare (where possible) completion rate, pass-rate and grade distribution between pandemic and pre-pandemic cohorts.

### 3.3 Survey Reflections

Responders reflected that the lack of in person education was detrimental in social and group interactions (engagement, feedback, discussion) fundamental to teaching and learning in all subjects (Table 4). Conversely certain positives were highlighted that may bring diverse strategies that enrich the learning and teaching experience. The benefits that were identified are varied and wider uptake make improve teaching and learning in biological NMR.

**Table 4:** Reflections of educators on the impact of the online learning environment.

| Positive Impact | Category | Negative Impact | Category |
| --- | --- | --- | --- |
| a combination of different methods<br>Being able to refer to recorded materials<br>easier for other teachers to join in exercises<br>For remote training only the student could type/mouse click, they did it all themselves.<br>H5P image ordering as a way of teaching triple resonance protein resonance assignment principle<br>it forced us to employ NMRbox to demonstrate various software packages and realized what a great resource it is<br>live demonstrations<br>making more time available<br>pre-recorded video allowed for more time for Q&A sessions<br>Quiz<br>screen sharing to identify difficulties<br>screen sharing when students did their assignments... is like sitting next to them and discussing with them what to do next<br>The use of NMRBOX.<br>Greater reach | facilitating<br>Sharing/ collaborating<br>Sharing/ collaborating<br>Practice<br>Learning<br>Practice<br>Practice<br>Facilitating<br>Facilitating<br>Assess<br>Practice<br>Practice<br>Facilitating<br>Sharing | (unable to do) ...experiments at the spectrometer<br>Demonstrations of use of NMR instrument was not as effective online as it was in person - the interactive nature of in-person demos of instrument use is hard to replicate online and most students required in person assistance even after demos were done online<br>difficult to have discussions<br>(no) Explanations at the white/black board<br>Getting feedback from the students.<br>Lectures themselves. It was really hard to keep students attention online.<br>(no) live demonstrations<br>No opportunity for the students to be trained at the magnet/console.<br>none<br>(no) Social contact<br>(lower) Student engagement<br>student participation was reduced<br>theory | Practice<br>Practice<br>Sharing<br>Sharing<br>Evaluating<br>Learning<br>Practice<br>Practice<br>n/a<br>Sharing<br>Sharing<br>Sharing<br>Learning |

Other feedback observed that the teaching landscape has changed again since the survey was designed (UK educators returning to face-to-face teaching) and as such follow-up for student cohort of 2022 would also be valuable information. Positive remote strategies included time-saving and ease-of-use activities as well as ways to implement practical experience. Negative remote strategies were primarily sharing/collaboration and a lack of opportunities for practical experience.

### 3.4 Focus Group

Focus group discussions (in person at conference where the preliminary results were discussed) and online with fellow educators in dedicated meetings highlighted a range of further insight. Working with pen and paper whilst sitting down with the students in groups was a major limitation in teaching new concepts in NMR to the students. The solution to demonstrate online and then send students away to work individually left educators with very little feedback as to how the students were meeting the problem-solving challenge of NMR and left the educatees without peer support or encouragement.

Recording materials (whilst not novel to recent years [Kwan *et al* 2019]) afforded a greater breadth of coverage to conceptually difficult topics and a wider range of teaching styles from multiple educators involved. Focus group discussions centred on greater prominence and sharing of such resources would be beneficial.

One perhaps surprising theme within the focus group was a lack of awareness of the length and breadth of online tools available for bioNMR education. This response may be due to the time-pressure NMR educators are under (and reliance on word-of-mouth to integrate new concepts and tools into teaching). A wider discussion and greater communication is one valuable outcome of this survey. NMR tools that may be most accessible are also potentially those most overlooked coming not from within biochemistry or chemistry education and instead within medicine. MRI harnesses the same quantum mechanical properties as NMR to image internal organs within the human body which of course requires medical communication between imaging specialists and patients/general public. The utility of NMR theory within MRI medical/analytical teaching [Westbrook 2017] may pave the way for clearer more tractable tools and teachings within patient education having a knock on effect for NMR.

## 4 Case Study

The N8 universities NMR group (ResoN8) delivered a remote workshop of NMR across many themes within this survey. In spring 2021 the ResoN8 NMR group delivered an all-online CCPN and ARIA workshop. This workshop hosted 27 participants – none of whom would be formally accredited to a degree programme most of whom were studying for a PhD in a directly related subject (Chemistry, biochemistry, biology etc.). The group found that benefits of team teaching [Govindarajan 2021] are different backgrounds (prior knowledge) of educators give different emphasis and multiple educators requires clear structure and objectives determined beforehand.

This workshop was possible due to the NMRBox.org platform – a free to use virtual environment with stable copies of CCPN Analysis (v2 and v3) and ARIA present. Using the ability to create a share group drives (and the invaluable support of the NMRbox team) all data, examples and outputs employed by the workshop were shared with all attendees without compatibility or versioning issues.

Reflecting on our practice of teaching via this virtual environment highlighted key benefits The major benefit of this mode of delivery was the virtual box environment which alleviated the pressures on the individual learner to install and set-up complicated software a major block to inclusion and psychological burden [Ravanelli *et al* 2014]. Possibly even more importantly, the platform afforded networking of the users accounts to provide a single set of data from all learners and educators to work from (not only saving download/upload of 5GB+ of files but also enabling the learners to work from ‘real’ NMR data). Dealing with the ‘process’ of raw data to structure is also a way of organising material (advanced organiser or scaffolding [Ausubel 1968]) and draws upon learners prior knowledge which in turn aimed to translate into meaningful learning such as that explored in Tian 2019 [Tian 2019].

Biological NMR in particular and STEM in general aligns extremely closely with inquiry- and discovery-driven teaching or evidence-based teaching [Petty 2009] which in turn is a constructivist [Piaget 1964] approach whereby probing ‘real’ data as a group can engage learners to reflect on the example in the context of their own interests and lens. Practice through the remote environment of NMRbox facilitated application of concepts in practice; a cognitivist approach providing additional opportunity to connect core concepts with deep learning.

## 5 Discussion

### 5.1 Limitations

The limitations of this survey are a very small sample size coupled with a diverse range of biological NMR applications and teaching strategies. Interpretation of questions across multiple educational structures and ambiguity in subject definitions. The study would benefit from a larger set of participants and potentially a more focussed scope. Even so from the modest sample size there was a range of responses sampling varying approaches to online teaching in biological NMR.

An emergency remote learning environment is not reflective of a planned remote learning environment. Motivation theory Maslow’s hierarchy of needs posits that deficiency needs unmet will impair the growth needs that underpin motivation. During enforced remote learning these deficiency needs may be unmet in a substantial proportion of learners (and educators) and as such motivation to engage with remote learning may be substantially lower than without the external pressures manifesting as a result of the pandemic. Educators may also have sought simplistic solutions and resorted to didactic learning more than would be expected in pre-pandemic teaching. Didactic learning aligns well with asynchronous teaching (recordings are by nature uni-directional and passive). This assumption is somewhat mitigated by survey questions that asked educators whether learners were also asked to use the software involved but too many participants responded that neither educator or learner engaged with online/software solutions to draw any strong conclusions in spite of most participants reporting that remote teaching was via a mixture of synchronous lectures (slides and voice), asynchronous lectures (slides and voice), polls, multiple choice quizzes etc.

### 5.2 Observations

The use of Topspin, CCPN and Pymol were most common among the respondents (between 61-76%). Over half of responders did not report use of online databases, despite multiple articles promoting them as educational resources [Laskowski et al 2017, Kinjo et al 2017, Zardecki et al 2016, Zardecki et al 2021]. Group work and engagement were negatively impacted by remote teaching. However, the flexibility of remote environment to share desktops (including spectrometer desktops) was positively received to engage with students and other teachers more effectively. Virtual computing environments (such as NMRbox) also afforded a stable platform for equitable access to multiple NMR packages in a stable environment while transferring requirement for high-performance computing from the individual to the service provider.

The Quality Assurance Agency for Higher Education [QAA] stipulates that masters/postgraduate level courses are built on knowledge established within undergraduate curriculum which can be an incorrect assumption due to the variety of routes into biological NMR. This means NMR theory/fundamentals may need direct signposting and revision of foundation principles. Further, problem solving requires confidence in content. Recorded and shared material can enhance and empower learners without the level of underpinning of their peers. This student-centred and progressivism approach [Kolb 2005] can be enhanced/ supported by the network of remote educators.

### 5.3 Recommendations

The need for more data and follow-up studies is apparent both within feedback from the survey (as higher education evolves post pandemic) and from the continued development of active-learning strategies and tools within the field of biological NMR.

The problem-solving behaviour of the cohort may also be a factor with students preference (or ability) to solve NMR problems conceptually or algorithmically [Domin & Bodner 2012] which would also impact the efficacy of teaching methods and resources used. NMR is conceptually difficult - considerations for Ausubels cognitive assimilation theory (CAT) – knowledge is assimilated from transferred from general to individual. Abstract concepts are tethered to personal projects.

While there is no one-size fits all solution to effective teaching and learning of biological NMR it is clear there are multitude of approaches UK (and global) educators employed during pandemic lockdowns and remote teaching.

Many suggested solutions may be limiting due to technology required – assuming students have the necessary equipment and CPU capabilities is a major drawback. Indeed, even when technology is available infrastructure within the learning environment such as stable internet access and sufficient power supply and/or battery life are all essential for success.

Low tech (mobile app) solutions to help work on problem-solving aspects (i.e. in lieu of face to face problem solving using pencil and paper) do not appear to exist in the field to date however given the explosion of online solution there may be suitable options that have yet to be applied to NMR.

Keep the remote/online classroom for at least a proportion of biological NMR teaching. Added value is that computational techniques and tools reinforce biochemistry theory and principals as well – a boon for learners and educators teaching final and post-graduate students where local knowledge and expertise may be limited and ability to share data remotely enhances network of educators and learners.

## 5.4 Acknowledgments

The author would like to thank CCPN annual conference (computational collaborative project for NMR) for hosting their flash presentation and poster of the preliminary findings to the Biological NMR community (20^th^ July 2022) and also CCPN, UKMRM (United Kingdom magnetic resonance managers) and NMRDG (Nuclear magnetic resonance discussion group) consortia for promoting the survey to their communities. The author is also indebted to ResoN8 NMR group [ResoN8] for discussions, feedback and evaluation.

## 6.2 Resources

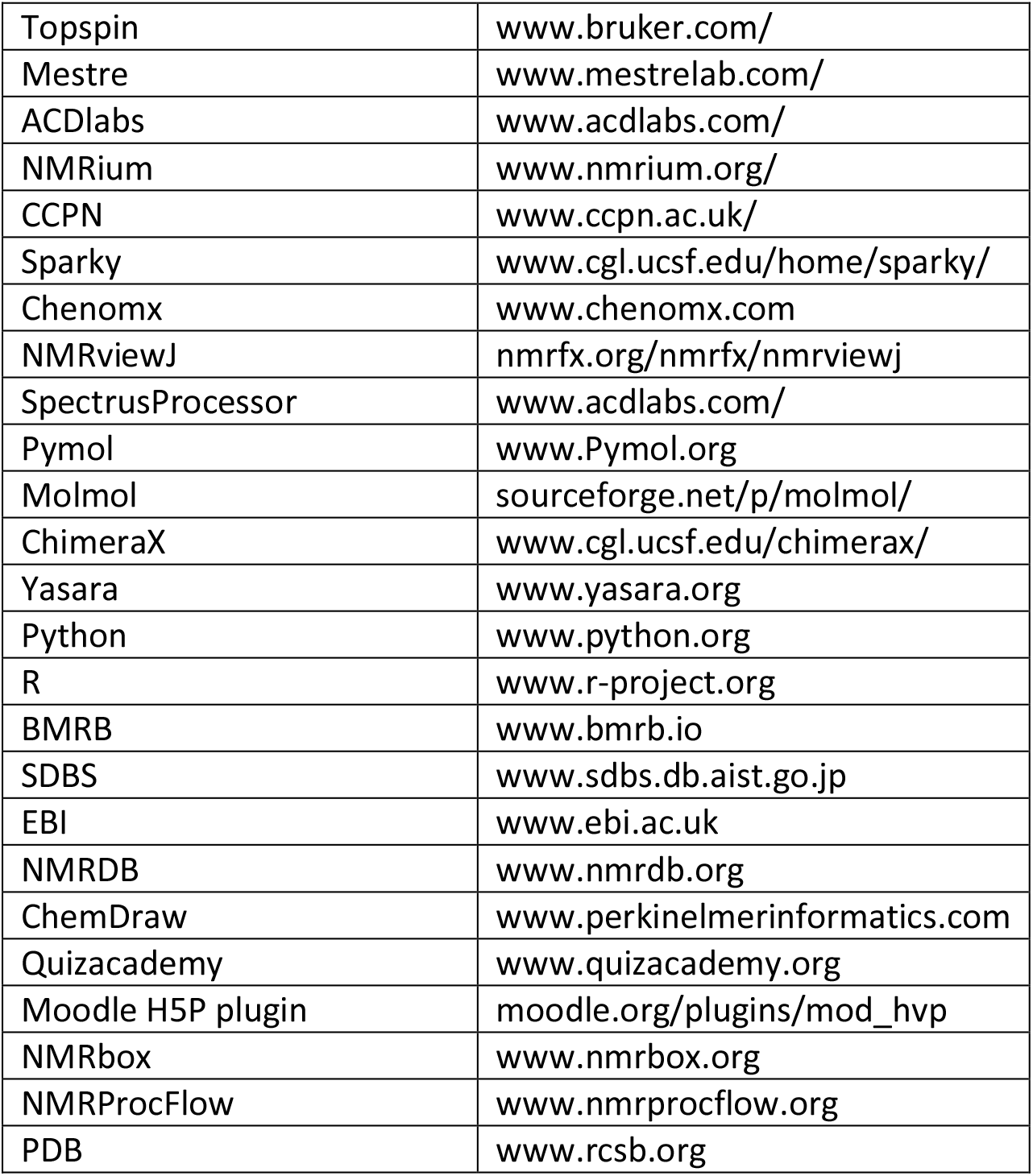

